# Delineation of the Trigeminal-Lateral Parabrachial-Central Amygdala Tract in Humans: An Ultra-High Field Diffusion MRI Study

**DOI:** 10.1101/2023.06.30.547270

**Authors:** Batu Kaya, Iacopo Cioffi, Massieh Moayedi

## Abstract

Orofacial pain is thought to be more unpleasant than pain elsewhere in the body due to the importance of the face in social, feeding, and exploratory behaviors. Nociceptive information from the orofacial region is carried to the brain via the trigeminal nerve (CNV) via the trigeminal brainstem sensory nuclear complex (VBSNC). Pre-clinical evidence revealed a monosynaptic circuit from CNV to the lateral parabrachial nucleus (latPB), which underlies the greater unpleasantness elicited by orofacial pain. The latPB further projects to the central amygdala (CeA), which contributes to the affective component of pain in rodents. However, this circuit has yet to be delineated in humans. Here, we aimed to resolve this circuit using 7T diffusion-weighted imaging from the Human Connectome Project (HCP). We performed probabilistic tractography in 80 participants to resolve the CNV-latPB-CeA circuit. The basolateral amygdala (BLAT) was used as a negative control, given that we did not anticipate CNV-latPB-BLAT connectivity. Connectivity strengths were compared using a repeated-measures ANOVA with factors ‘hemisphere’ (left; right), and ‘target’ (CeA; BLAT), with sex included in the model for both pilot and validation samples. Only the ‘target’ factor was significant in both samples (*F*_Pilot_ = 11.4804, *p* = 0.005; *F*_Validation_ = 69.113, p < .001). *Post hoc* tests showed that the CeA had significantly stronger connectivity strength than the BLAT (*p_Tukey-Pilot_* = 0.005; *p_Tukey-Validation_* < 0.001). □This study delineates the human CNV-latPB-CeA circuit for the first time *in vivo.* This circuit may provide a neuroanatomical substrate for the affective dimensions of orofacial pain.

**SUMMARY:** This study delineates the human trigeminal-parabrachio-amygdalar circuit *in vivo.* This circuit may provide a neuroanatomical substrate for the affective dimension of orofacial pain.

## INTRODUCTION

Somatosensory information, including tactile, noxious, and thermal inputs from the orofacial region, is carried to the central nervous system by the trigeminal nerve (CNV). Afferents from the trigeminal ganglion synapse onto the trigeminal brainstem sensory nuclear complex (VBSNC), and further relay to other brain regions [27]. The unique neural architecture of the orofacial region is thought to undergird the evolutionary, emotional, and psychological primacy of information coming from the face [16,49,52]. Indeed, experiments have shown that participants display increased fear of orofacial pain compared to pain in limbs despite similar stimulus intensities [50]. Additionally, experimental facial pain associated recall—as opposed to experimental hand pain associated recall—correlates with increased activity in memory-related regions (e.g., hippocampus) [49]. However, the mechanisms underlying the amplification orofacial pain associated negative affect remain unclear.

The affective dimension of pain comprises constructs such as fear of pain, suffering and pain distancing, i.e., nocifensive behaviors [38,43]. The parabrachial nucleus, located in the brainstem, is an integrative hub for autonomic, nociceptive and multisensory inputs and is involved in orchestrating motor outputs in response to noxious stimuli [14]. Second order trigeminal afferents synapse onto the lateral division of the parabrachial nucleus (latPB) via branches of the trigeminothalamic tract. Further, this region has extensive connections to pain modulatory regions, such as the central nucleus of the amygdala (CeA), the periaqueductal gray (PAG), and the rostral ventromedial medulla (RVM) [21,40,41].

In a recent study, Rodriguez et al [44] discovered a novel circuit in mice between latPB and CeA that elicited increased pain-associated distress behaviors in response to noxious stimuli to the whisker pad, compared to the hind paw. Strikingly, this direct CNV ganglion-latPB-CeA circuit bypasses the canonical VBSNC processing pathway, and thus, is thought to contribute directly to the affective dimension of pain, given the dense connectivity of the latPB and the CeA. We posit that this circuit may underlie the heightened unpleasantness associated with orofacial pain, however, no studies to date have investigated the presence of this pathway in humans.

Subdivisions of brainstem nuclei (e.g., latPB) and of subcortical brain regions (e.g., CeA) are becoming increasingly more accessible due to the proliferation of probabilistic atlases [13,53,56,57]. Open-source high field diffusion weighted imaging (DWI) data from databases such as the Human Connectome Project (HCP) [59] make possible the use of tractography [5] to translate trigeminal neural circuitry identified in rodents to humans. Here, we aim to delineate the CNV-latPB-CeA circuit *in vivo* in an exploratory and a validation sample using ultra high-field (7T) imaging data from the HCP database. As a secondary aim, we will investigate whether there are sex differences in the connectivity strength of this circuit.

## METHODS

All procedures were approved by the University of Toronto’s Human Research Ethics board (Protocol Number: 40458). The data that support the findings of this study are openly available in the HCP database at https://db.humanconnectome.org, HCP S1200 Release (February 2017).

### Participants

We delineate the CNV-latPB-CeA circuit in two independent samples (a pilot cohort, and a validation cohort) from the HCP database’s S1200 release, which includes preprocessed 7T diffusion images for 184 healthy young adults and is the final release from the Washington University and University of Minnesota consortium. Participant inclusion/exclusion criteria for recruitment is publicly available [59]. Briefly, exclusion criteria included significant history of psychiatric disorders, substance abuse, neurological or cardiovascular disease, epilepsy, any genetic disorder, neurodegenerative disease, traumatic brain injury, ongoing chemo-and/or immunotherapy, and MRI contraindications. Readers interested in the full exclusion criteria are encouraged to refer to Supplemental Table 1 in Van Essen et al. (2013) [59].

In this study, the pilot sample comprised of 15 subjects (8 females, 7 males) randomly selected from the S1200 release. Note that the HCP does not provide ages for participants, but simply age ranges (22-25, 26-30, 31-35 years of age). To assess differences between the circuit of interest and a control circuit, a repeated measures analysis of variance (ANOVA) was performed with two factors (hemisphere; target) and sex as a between-subjects factor. Please see section *Region of Interest Definition: Control Circuit* for more details. The data were natural logarithm (ln)- normalized due to positive skew in both samples. Normal distribution was confirmed for data points after transformation via Q-Q plots and Shapiro-Wilk tests. The sex-by-target interaction from the pilot cohort was used to obtain an effect size for power calculation. The sample size for the validation group was calculated using the G*Power software [19], using a partial-eta-squared (η_p_^2^) = 0.043 (equivalent to an effect size *f* = 0.212). The power calculation revealed that 32 participants are required for adequate power to reproduce the analysis in an independent sample (*a* = 0.5, power (1-ß) = 0.80). A total of 80 participants (28 males, 52 females) were randomly selected from the HCP database to account for potential outliers, and to allow for sex disaggregated analyses. Again, we note the HCP does not provide ages for participants, but simply age ranges (22-25, 26-30, 31-35 years of age). There was no overlap between the samples.

### HCP Imaging Parameters

All participants in this study underwent MRI scanning in a 3T and a 7T scanner. The 3T scans were acquired at the Washington University in St. Louis, using a customized Siemens 3T “Connectome Skyra,” with a standard 32-channel head coil and a body transmission coil. Participants were then flown to the Center for Magnetic Resonance, University of Minnesota for a 7T scans acquired with a Siemens Magnetom scanner using a Nova32 32-channel Siemens receive head coil with an incorporated head-only transmit coil that surrounds the receive coil from Nova Medical.

Whole brain structural T1 scans were acquired at 3T with the following parameters: repetition time (TR) = 2400□ms, echo time (TE) = 2.14□ms, field of view (FOV) = 224 ×□224, and 0.7 mm isotropic voxels. The spin-echo EPI diffusion-weighted scans were collected at 7T over four runs with the following parameters: TR = 7000□ms, TE = 71.2□ms, FOV = 210 ×□210, 1.05□mm isotropic voxels, and 132 slices. These were acquired with two sets of gradient tables, each with two different *b*-values (b= 1000 s/mm and b = 2000 s/mm) interspersed with an equal number of acquisitions on each shell within each run. The uniform distribution of directions across shells was ensured with the Emmanuel Caruyer toolbox [12]. Each set included 65 diffusion-weighted directions plus 6 non-diffusion weighting images (B0s) interspersed across each run. Two of four acquisitions were performed once with each anterior-to-posterior and the other two with posterior-to-anterior encoding polarities. The 1200 Subjects Data Release Reference Manual by HCP can be consulted for further details for data acquisition (https://www.humanconnectome.org/storage/app/media/documentation/s1200/HCP_S1200_Rele ase_Reference_Manual.pdf.).

### MRI and Statistical Analysis

#### HCP Preprocessing

For each participant, we downloaded the preprocessed whole brain T1 scan (T1w_acpc_dc_restore image) and the 7T diffusion weighted scan. The HCP pre-processing pipeline scripts are publicly available on Github (https://github.com/Washington-University/HCPpipelines). Readers are encouraged to consult the HCP Consortium’s article on the minimal preprocessing pipelines for a thorough review [22].

Briefly, structural runs were gradient distortion corrected, and then aligned using a rigid-body alignment with 6 degrees-of-freedom (DOF) and averaged to create one T1 image if there were more than a single run. The FMRIB’s software library (FSL, v 5.0.6; https://fsl.fmrib.ox.ac.uk/fsl) Linear Registration Toolbox (FLIRT) [29,30,55] was used to register the resultant image to the Montreal Neurological Institutes (MNI) standardized space template using 12 DOF. After AC-PC alignment, FSL’s brain extraction tool (BET) [31] was used to skull strip the structural T1 images. Finally, readout distortion correction was applied to generate the final T1 image.

For diffusion weighted images, FSL was used to normalize B0 intensity using FSL’s Diffusion toolbox (FDT) [30], after which susceptibility induced B0 field deviations were calculated. The eddy tool [3] was used to model eddy current distortion and subject motion. After correcting for these deviations, the diffusion weighted image was resampled into subject’s native space using FLIRT. A binary brain mask was generated for fibre orientation estimation using FSL’s BEDPOSTx tool [5,6].

### Study-Specific Preprocessing

Diffusion weighted and T1 images were skull stripped. For DWI scans, we aimed for liberal skull-stripping to ensure the inclusion of CNV fibers in the resultant brain mask. For this purpose, BET (v6.0.3) [31] was used to remove non-brain tissues from the non-diffusion-weighted (B0) image in the DWI images. Each skull stripped B0 scan was quality controlled (QC) to ensure the inclusion of CNV fibers. Voxels were manually drawn and added to the brain mask as necessary to ensure full capture of the CNV. The whole brain T1-weighted scans were skull stripped using the SynthStrip toolbox [26] in FreeSurfer v7.3.2 [20]. Affine registration with FLIRT [29] was performed in the FSL diffusion toolbox (FDT) for each participant’s diffusion image to the T1-image, and further to the 2mm MNI152 template. Additionally, FSL’s non-linear registration tool (FNIRT; [2]) was run using FSL’s T1_2_MNI152_2mm configuration file to generate a warp matrix that was later used in constructing group tractograms for the circuit of interest in the study. FNIRT matrices were preferred over FLIRT ones for group tractograms due to reduced distortions and improved brainstem registration. Matrices from the FLIRT registration were used to transform the probabilistic seeds for CeA, BLAT, PAG, and latPB in MNI space to diffusion space for each subject and were visually inspected for accuracy. A tensor was fit using FSL’s DTIFIT to visualize major fibre orientations for the entire brain. CNV seeds and exclusion masks were drawn manually. Please see *MRI and Statistical Analysis: Probabilistic Tractography* for more details. Two principal fiber directions were modeled for every voxel using BEDPOSTx. Lastly, we used Probtrackx [5] to assess connectivity strength between our regions of interest. The resulting tracts maps were resampled using the non-linear registration matrices from diffusion space to MNI space for visualization purposes, as previous work has shown that FNIRT resampling can improve visualization of white matter tracts [62]. Please refer to *Supplementary Figure 1* for a summary flowchart of the study-specific preprocessing pipeline.

### Regions of Interest Definition

Figure 1 provides a schematic of the regions of interest (ROIs) used in the study.

**Figure 1:**
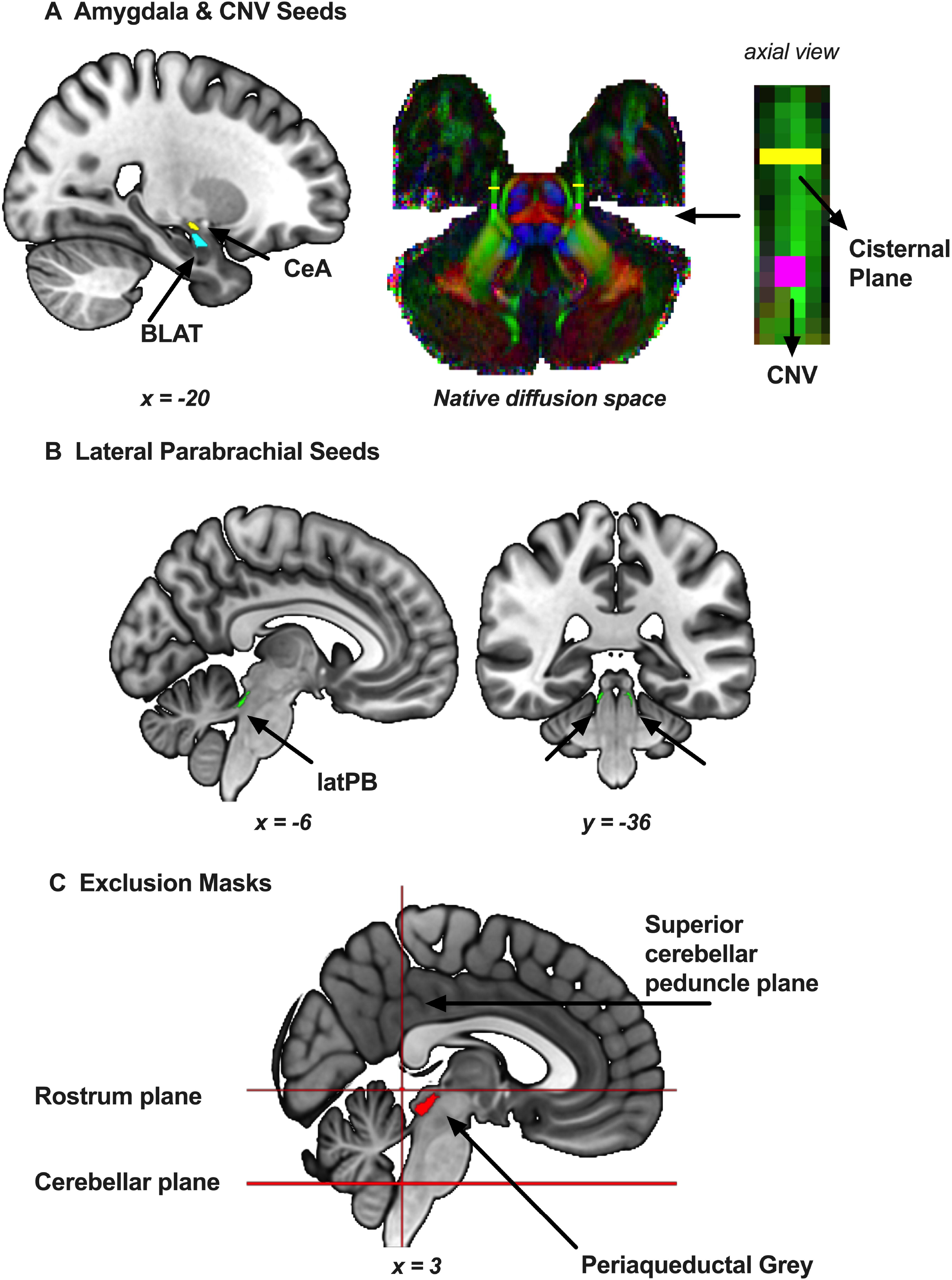
Seeds and exclusion masks used in tractography. CeA: Central amygdala, BLAT: Basolateral amygdala, CNV: Cranial nerve 5 (Trigeminal nerve), latPB: Lateral parabrachial nucleus. MNI coordinates can be found below each brain slice.

#### Cranial Nerve V

The cisternal segment and the root entry zone of the CNV were used as a proxy for the trigeminal ganglion to capture trigeminal afferents coursing to the central nervous system. Recent work shows that a two ROI method is superior to single and three ROI strategies for tracking the trigeminal nerve [61]. Additionally, the cisternal segment was used as a waypoint to ensure that samples were propagated toward the brainstem. The first eigenvector image produced by the tensor fit was modulated by fractional anisotropy values to visualize fiber direction with color. The CNV was then located in the axial plane. A 2×2×2 cube (8 voxels in total with a volume of 9.261 mm^3^) was drawn at the CNV root-entry zone. Then, a 1×4×4 cisternal segment was drawn anterior to the CNV ROI (16 voxels in total with a volume of 18.522 mm^3^). We located the widest fanning segment of CNV fibers anterior to the REZ and outside the temporal lobes when picking the location of the cisternal segment. This process was repeated bilaterally.

#### Amygdala

The Tyszka and Pauli atlas provides probabilistic ROIs for amygdalar subnuclei in standard MNI space [57]. This atlas is based on a sample of 168 healthy adults with an age range of 22-35 years from the HCP database. Two seeds, CeA and basolateral amygdala (BLAT), were used in probabilistic tractography as final targets and the origin when propagating putative tracts in an ascending, (i.e., brainstem to subcortex) and descending manner (i.e., subcortex to brainstem), respectively. Several naming conventions exist for the amygdalar basolateral complex [35]. To avoid confusion, we followed the classifications used in this probabilistic atlas. The BLAT seed in this study does not include the lateral and accessory basal/basomedial subnuclei. Additionally, we chose to remove the parvocellular division of the basolateral nucleus from the BLAT seed, since the developers of the atlas noted that this specific nucleus was merged into an unresolved nucleus (paralaminar) when creating the final seeds for their probabilistic atlas. In short, the BLAT seed in this study included the fully defined dorsal and intermediate divisions of the basolateral nucleus.

#### Lateral Parabrachial Nucleus & Periaqueductal Gray

The Brainstem Navigator atlas (v0.9; https://www.nitrc.org/projects/brainstemnavig/) provides probabilistic seeds for brainstem subnuclei crucial for sensorimotor, arousal and autonomic functions [8]. Semi-automatic and manual segmentations of these nuclei were carried out using 7T, multi-contrast MRI data from *in vivo* to generate the seeds. In this study, the 35% thresholded latPB seeds were used as waypoints in probabilistic tractography. The probabilistic periaqueductal gray (PAG) seed was thresholded at 50% to generate an exclusion mask. The difference in threshold values between the seeds is due to their utility in the study: the latPB is used as a target, and thus we were more lenient to ensure we captured individual variability in the seed, whereas the PAG is used as an exclusion mask, and so we were more conservative to ensure we do not exclude tracts of interest. The specific atlas labels for the latPB seeds were LPB-l and LPB-r, and PAG for the PAG seed. Both the amygdalar and brainstem seeds were binarized.

### Probabilistic Tractography

Diffusivity characteristics of water in brain tissue can be modeled to study white matter tracts *in vivo* [4]. Probabilistic tractography generates a tractogram quantifying probable structural connectivity. From a voxel, a single sample in one of the principal diffusion directions is initiated, and then the streamline is propagated posteriorly. This process is repeated for a specific number of samples to calculate a ratio of the number of propagated streamlines intersecting with the dominant path distribution [5]. The resultant probabilistic path distribution map thus provides a quantitative measurement of putative circuits in the brain.

In this study, we performed probabilistic tractography to assess CNV-latPB-CeA connectivity. The CNV-latPB-BLAT served as a control circuit since CeA is the only amygdalar subnucleus with dense connectivity with the brainstem [37]. Each subject had a total of 8 tractograms: Within one hemisphere, fibers were constructed in both directions (i.e., Origin-to-Target; Target-to-Origin) for each circuit to account for directional biases during acquisition [58] and fiber fanning [32]. This resulted in four tractograms per hemisphere. We used the modified Euler algorithm and set the number of samples to 10,000. A curvature threshold of 0.1 was used to allow for sharper turns for paths crossing from brainstem to subcortical regions. Path distribution was corrected for the length of the pathways.

Tract propagation was further specified using exclusion planes to limit the boundaries of the path distribution space in the brain, and by incorporating waypoints as relays that had to be crossed prior to reaching target. More specifically:

1. A midline mask was drawn in diffusion space to constrain tracts to a single hemisphere.
2. An axial plane below the rostrum of corpus callosum was drawn in diffusion space to discard projections from CeA to the neocortex.
3. A coronal plane across the superior cerebellar peduncle was drawn in diffusion space to discard cerebellar fibers. We ensured that there was no contiguity between the latPB seeds and the exclusion plane for each subject.
4. An axial pontine plane was drawn above the medulla oblongata in diffusion space to remove potential pyramidal and spinothalamic fibers.
5. Due to PAG’s extensive projections to the latPB [34], we added the PAG seed as an exclusion seed while propagating tracts.
6. Additionally, the probabilistic mask of the CNV motor nucleus from a previous study [33] was thresholded at 50%, and used as exclusion mask to discard pontine CNV fibers [23].
7. To ensure that tracts did not propagate past CeA or CNV depending on the direction, we set the target seed both as a waypoint and a termination mask.

Additionally, CS and latPB seeds were included as waypoint mask independent of directionality. Ascending tracts were propagated from CNV towards CP, then latPB, and eventually to their amygdalar target. Tracts in the opposing direction originated in the amygdala and terminated at CNV via latPB and CS, in this order.

Connectivity strengths, based on tractograms delineated in opposing directions for each circuit, were corrected for seed size, averaged and thresholded to generate a single tractogram per circuit (Please see Section: *Statistical Analyses*). In total, four final tractograms were generated: Right CNV-latPB-CeA, Left CNV-latPB-CeA; Right CNV-latPB-BLAT and Left CNV-latPB-BLAT.

### Statistical Analyses

We quantified connectivity strength for the circuits in the study using the “waytotal count”, which is the number of streamlines that reach the final target seed in accordance with pre-defined user criteria (See section *Probabilistic Tractography* for more details). Initially, the total number of streamlines is equal to the product of the number of voxels in the seed of origin and the number of total streamlines to be used in propagation, which was set to 10,000 in this study. However, given the differences between the number of voxels between the trigeminal and amygdalar seeds, we normalized the number of streamlines that reached the target by dividing the count by the number of voxels in the initial seed. Then, normalized waytotal counts between the same origin and target seeds (e.g., Right CNV to Right CeA, Right CeA to Right CNV) were averaged. In the end, there were four probabilistic connectivity strength measurements per participant: Right CNV-latPB-CeA, Right CNV-latPB-BLAT, Left CNV-latPB-CeA, and Left CNV-latPB-BLAT. We further thresholded these tracts at 2% to avoid spurious connections. Q-Q plots, and Shapiro-Wilk tests were run to assess normality visually, and statistically with a Bonferroni-corrected alpha of *p* < 0.0125 (See *Results* for more details). Given that data were not normally distributed, we natural log (ln)-normalized the data to run a repeated measures analysis of variance (rmANOVA) to assess within and between group differences. Sex was added as a between-subjects factor explore potential sex differences in the connectivity strength of the circuit of interest. *Post hoc* tests were performed using Tukey’s tests with significance set at *p* < 0.05. In the pilot sample, ln-normalization resulted in no loss of data points, and the test were run in all 15 participants. In the validation sample, the final sample size was equal to 67 after ln normalization, as some data could not be normalized (note that this exceeds the required sample of 32 individuals, and 64 individuals for disaggregated analyses).

## RESULTS

### Participants

The final pilot sample included 15 participants (7 males, 8 females; 27.4 ± 1.5 years, 31.8 ± 2.3, respectively) and the final validation sample included 67 participants (22 males, 45 females; 28.1 ± 3.8, 30.1 ± 2.7, respectively).

### Normality Testing

Shapiro-Wilk’s tests were run to assess normality of tractograms. In the pilot sample, all four connectivity strengths diverged significantly from normal distribution (Right CNV-latPB-CeA: *S-W*_15_ = 0.791, *p* = 0.003; Right CNV-latPB-BLAT: *S-W*_15_ = 0.560, *p* < 0.001; Left CNV-latPB- CeA: *S-W*_15_ = 0.410, *p* < 0.001 and left CNV-latPB-BLAT: *S-W*_15_ = 0.753, *p* < 0.001). After natural logarithmic normalization, all tracts had normal distribution (Right CNV-latPB-CeA: *S-W*_15_ = 0.956, *p* = 0.615; Right CNV-latPB-BLAT: *S-W*_15_ = 0.952, *p* = 0.550; Left CNV-latPB- CeA: *S-W*_15_ = 0.894, *p* = 0.078 and left CNV-latPB-BLAT: *S-W*_15_ = 0.919, *p* = 0.187).

Kolmogorov-Smirnov tests were run to assess the normality of tractograms in the validation sample since the sample contained over 50 subjects. All connectivity strengths from the validation sample were non-normally distributed (Right CNV-latPB-CeA: *K-S*_80_ = 0.192, *p* < 0.001; Right CNV-latPB-BLAT: *K-S*_80_ = 0.374, *p* < 0.001; Left CNV-latPB-CeA: *K-S*_80_ = 0.314, *p* < 0.001 and left CNV-latPB-BLAT: *K-S*_80_ = 0.314, *p* < 0.001). Data distribution was normal after ln-normalization (Right CNV-latPB-CeA: *K-S*_67_ = 0.071, *p* > 0.05; Right CNV-latPB- BLAT: *K-S*_67_ = 0.057, p > 0.05; Left CNV-latPB-CeA: *K-S*_67_ = 0.083, *p* > 0.05 and left CNV- latPB-BLAT: *K-S*_67_ = 0.093, *p* > 0.05).

### Connectivity Strengths of CNV-latPB-CeA Circuits and Sex Differences in the Circuit

A rmANOVA was carried out for the pilot sample using the natural log normalized data. Two factors were included: target (CeA vs. BLAT) and hemisphere (Right vs. Left), with sex added as a between-subjects factor in the model. The sphericity assumption was met since the model had only two levels. Levene’s homogeneity of variances test was non-significant for all four circuits with *p* > 0.05. Overall, only the target factor was significant (*F_1,13_* = 11.4804, *p* = 0.005, *η^2^_p_* = 0.469). A post-hoc t-test with the Tukey correction revealed a significant difference for the target factor, where CeA is significantly higher than BLAT (*t* = 3.39 and *p_Tukey_* = 0.005; see *Figure 2*). Potential sex differences in the connectivity strength of the circuits were explored by looking at three different interactions between factors: 1. hemisphere × sex, 2. target × sex, 3. hemisphere × target × sex. None of these interactions were significant in the pilot sample analyses (Target ×sex: *F_1,13_* = 0.5896, *p* = 0.456, *η^2^_p_* = 0.043; Hemisphere × sex: *F_1,13_* = 0.0332, *p* = 0.858, *η^2^_p_* = 0.003; Hemisphere × target × sex: *F_1,13_* = 0.8918, *p* = 0.362, *η^2^_p_* = 0.064.). There were no between-subjects effect for ‘sex’ (*F_1,13_* = 0.0163, *p* = 0.900, *η^2^ _p_* = 0.001).

**Figure 2:**
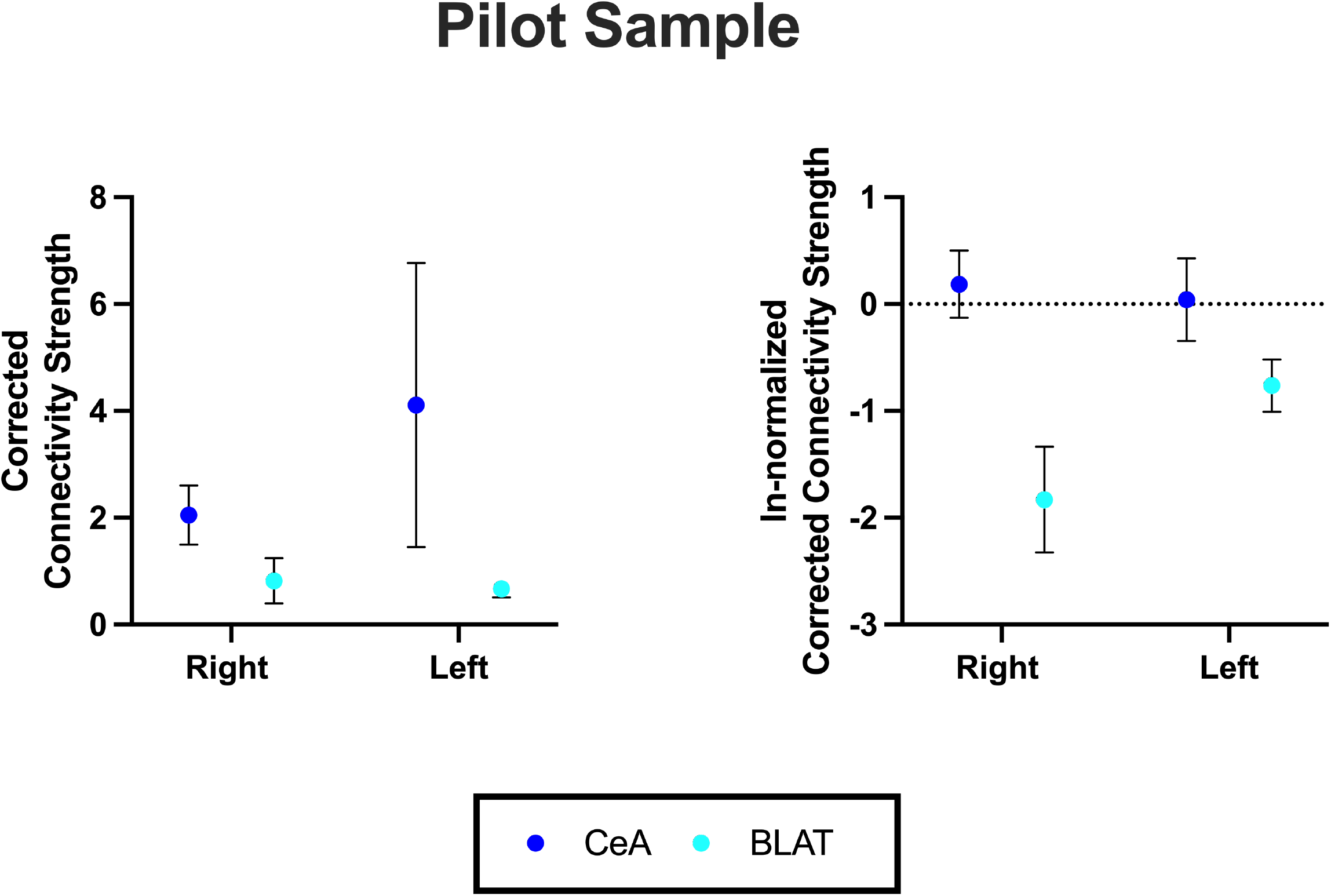
The CNV-latPB-CeA circuit has stronger connectivity than the CNV-latPB- BLAT circuit in the pilot cohort. Positively skewed (left) and ln-normalized (right) group means (±SEM) are shown for the pilot sample. After ln-normalization, a repeated measures ANOVA with sex as a between-subjects factor was significant for the target ‘factor’ (*F*_1,13_ = 11.4804, *p* = 0.005, *η^2^_p_* = 0.469).

We replicated these findings in our validation sample. The main effect of the factor ‘target’ was significant (*F_1,65_* = 69.113, *p* < 0.001, *η^2^_p_* = 0.515). A post-hoc t-test with the Tukey correction revealed a significantly stronger connectivity strength for the CeA than the BLAT (*t* = 8.31, *p_Tukey_* < 0.001; see *Figure 3*). Bilateral group tractograms for the CNV-latPB-CeA circuit were created using FLIRT and FNIRT matrices respectively (see *Figure 4*). None of the interaction terms that included sex were significant (Hemisphere × sex: *F_1,65_* = 0.221, *p* = 0.640, *η^2^_p_* = 0.003; Target × sex: *F_1,65_* = 1.707, *p* = 0.196, *η^2^_p_* = 0.026; Hemisphere × target × sex: *F* = 1.795, *p* = 0.185, *η^2^_p_* = 0.027). There were no between-subjects effect for ‘sex’ (*F* = 1.55, *p* = 0.218, *η^2^_p_* = 0.023).

**Figure 3:**
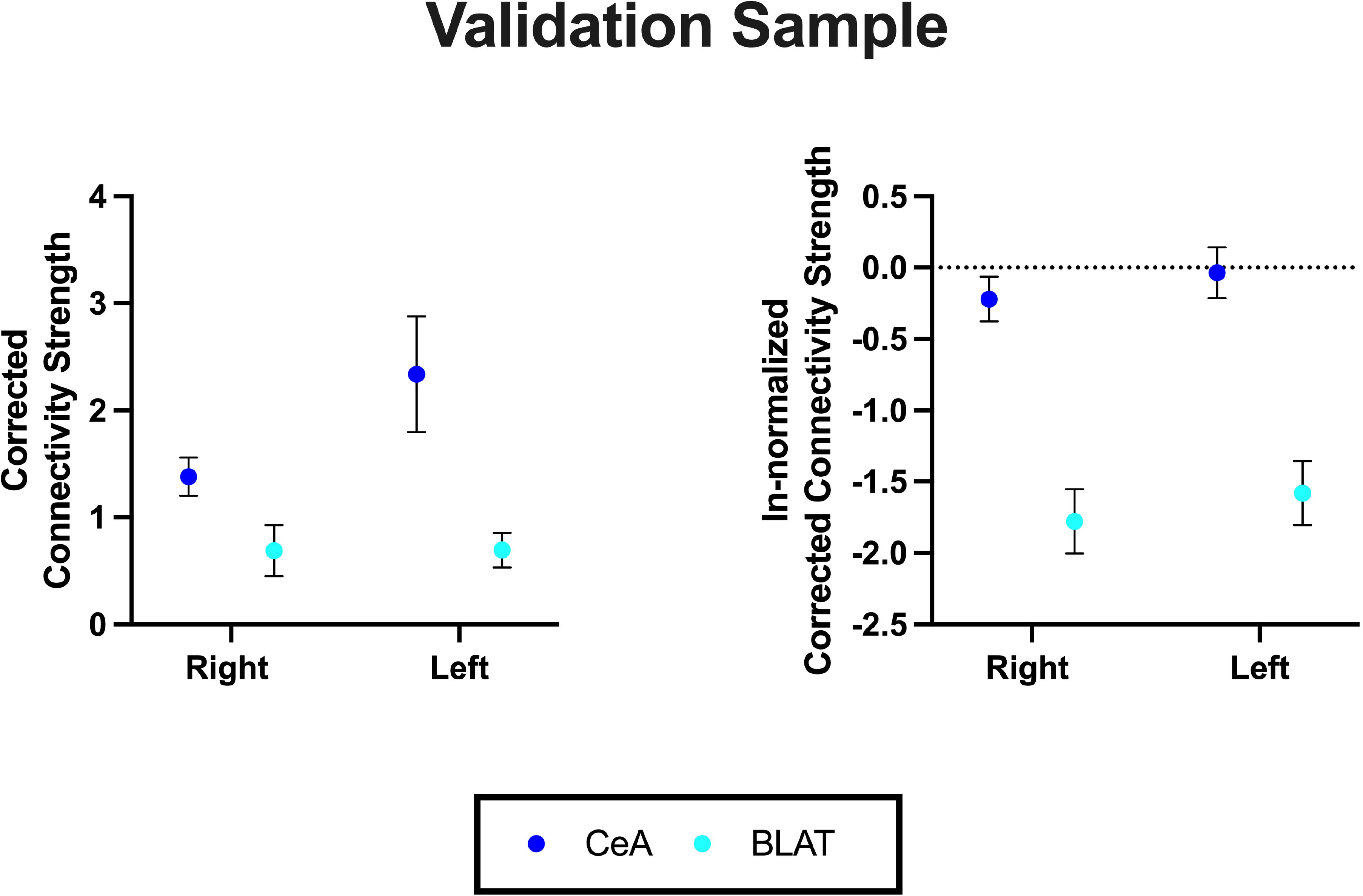
The CNV-latPB-CeA circuit has stronger connectivity than the CNV-latPB- BLAT circuit in the validation cohort. Positively skewed (left) and ln normalized (right) group means (±SEM) are shown for the validation sample. After ln-normalization, a repeated measures ANOVA with sex in the model was significant for the factor ‘target’ (*F_1,65_* = 69.113, *p* < 0.001, *η^2^_p_* = 0.515). Note that there were no significant interaction effects for hemisphere × sex: *F_1,65_* = 0.221, *p* = 0.640, *η^2^_p_* = 0.003; target × sex: *F* = 1.707, *p* = 0.196, *η^2^_p_* = 0.026; or hemisphere × target × sex: *F_1,65_* = 1.795, *p* = 0.185, *η^2^_p_* = 0.027). There were no significant between-subjects effect for ‘sex’ (*F_1,65_*= 1.55, *p* = 0.218, *η^2^_p_* = 0.023).

**Figure 4:**
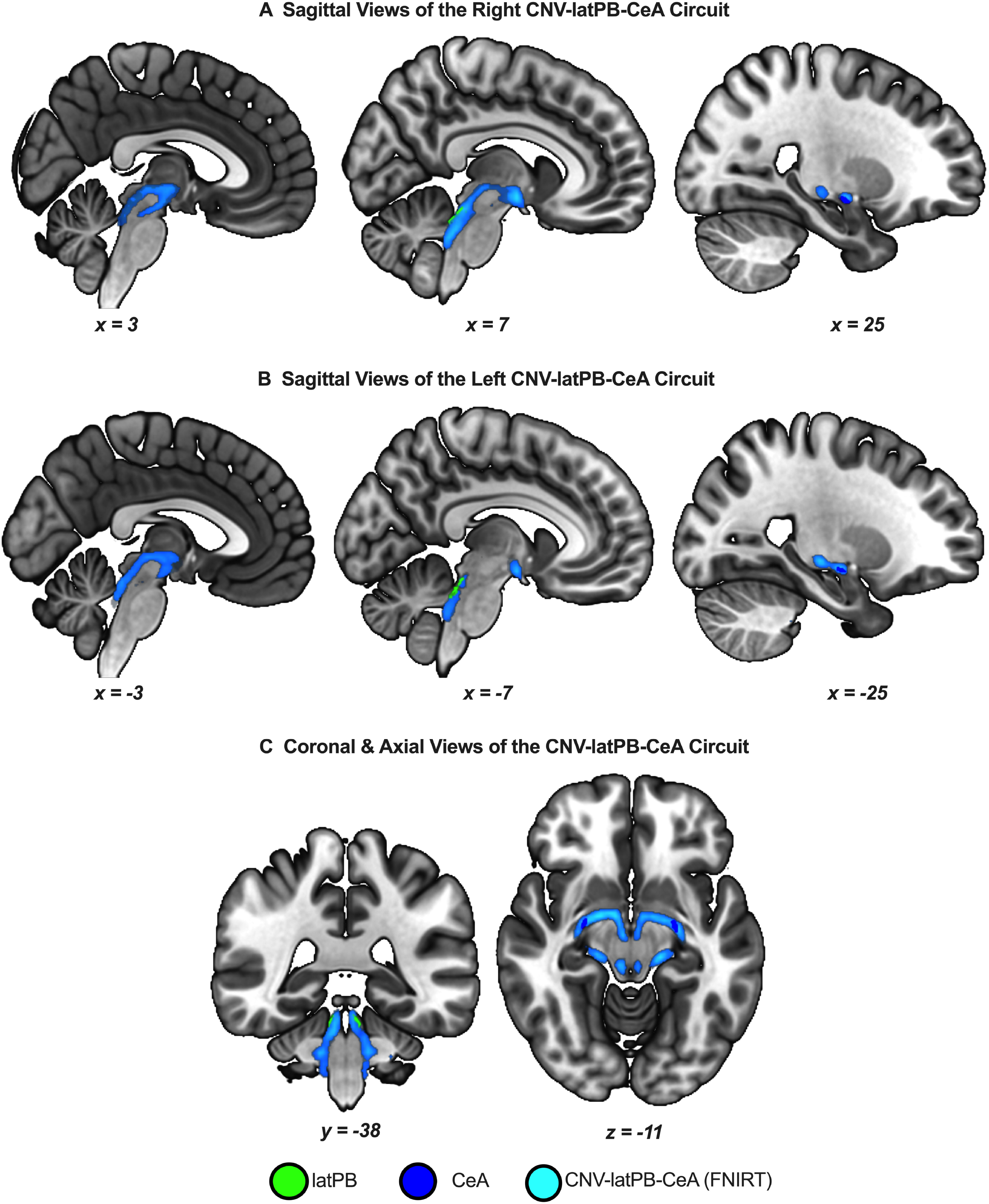
The CNV-latPB-CeA can be delineated *in vivo* using ultra high field 7T images. Group tractograms are based on N=80, thresholded to show at least 35% overlap across subjects in the validation sample. Panels A & B show the right and left circuits, respectively. The clear termination in the CeA, and the omission of potential PAG relays can be seen in the figure. Panel C illustrates passage of the tracts through the lateral parabrachial nuclei in concordance with the waypoint specification in the probabilistic tractography algorithm. The group tractograms were created using the linear FNIRT algorithm.

We then performed independent samples t-tests to explore our sample in a sex-disaggregated manner using the tractograms that survived the ln-normalization (45 females, 22 males). None of the tractograms revealed sex-specific differences in probabilistic connectivity strength in our validation sample (all *p > 0.05*).

## DISCUSSION (1297/1500)

In this study, we successfully delineated for the first time a direct pathway originating in the trigeminal root entry zone and terminating in the central nucleus of the amygdala via a relay in the lateral parabrachial nucleus (latPB) *in vivo* in two independent young adult samples. We exploited significant increases in image resolution in ultra-high field (UHF) 7T DWI from the HCP to counteract constraints inherent to brainstem imaging and probabilistic tractography such as weak contrast between white and grey matter [51], and fiber fanning [32], respectively. Additionally, we incorporated seeds from two recent probabilistic atlases, Tyszka-Pauli atlas, and the Brainstem Navigator, that were created using UHF 7T images and expert manual delineations for the ROIs. This proof-of-concept study highlights the possibility of investigating the functional role of this circuit in the affective dimension of pain in humans.

The affective dimension of pain is undergirded by specific neural circuits that regulate behavioral, emotional, and autonomic responses to noxious stimuli. The amygdala is a medial temporal brain region responsible for assigning an emotional value to environmental stressors [35], amongst other functions. Specifically, in the context of pain, the functional role of the amygdala is thought to be related to enhanced suffering, fear of pain and initiating motivational drive to mount nocifensive behaviours [40]. The CeA projects to several brainstem nuclei to modulate autonomic and motor responses to stimuli [42], and is comprised of three main subnuclei: the central medial, lateral central, and capsular central amygdala [60]. The capsular and lateral subnuclei of CeA are dubbed the “nociceptive amygdala” due to the direct excitatory inputs from the PB [1]. Similar to the amygdala, the PB plays an integrative role in modulating behavioral outputs in response to internal and external stressors [14]. Specifically, in rodents, the latPB receives polymodal nociceptive input from lamina 1 neurons in the dorsal horn [15,25]. In comparison, orofacial nociceptive information gets transmitted via the VBSNC. The VBSNC is a nuclear complex comprising the trigeminal main sensory nucleus, as well as the spinal trigeminal nucleus, which is made up of three subnuclei: oralis, interpolaris, and caudalis [52]. The entire complex receives noxious input from the periphery, but caudalis has two overlapping somatotopic maps of the orofacial region [17]. This organization is unique to the orofacial region and contributes to the interpretation of the orofacial region having important psychosocial function. Second-order afferents emerge from the VBSNC and further to other brain regions. Although spinal and trigeminal nociceptive input overlap partially in PB, in latPB these show anatomically segregated projection distributions [54], suggesting that bodily and orofacial pain are processed differentially—much like in the ventrolateral thalamus. A robust body of tracing studies performed in rats show that the CeA and the PB are connected via the spino-trigemino-parabrachial circuit [7,28,36]. However, the neural circuity undergirding enhanced affective responses to orofacial pain—as opposed to bodily pain—remains an outstanding question.

Rodriguez et al. [44] identified a novel direct monosynaptic circuit from the trigeminal ganglion to the external latPB subnucleus, which bypasses the VBSNC in mice. Three independent assessments of increased affect in response to the optogenetic activation of this circuit were carried out. Awake behaving mice emitted increased distress vocalizations, engaged in an aversive memory driven choice in a conditioned place aversion assay, and showed robust avoidance behaviors under optogenetic activation. Another study showed that external latPB- CeA connectivity underlies the formation of aversive memory during a Pavlovian threat conditioning paradigm involving electric shocks in context and a cue-dependent manner [24]. Mice with the presynaptic functional inactivation of PB-CeA displayed intact responses to the sensory-discriminative components of the noxious stimuli, i.e., hind paw lick latency, yet had an ablation of escape jumping behavior (i.e., nocifensive behaviours). These results suggest that spino-parabrachio-amygdaloid pathway modulates affective and not sensory-discriminative responses to noxious stimuli in mice.

Few studies have investigated whether and how orofacial nociceptive input is processed differently than spinal nociceptive input in humans. Sambo et al. [46] conducted two studies on the hand blink reflex (HBR) to characterize the properties of the defensive peripersonal space surrounding the face. They first showed that the waveform characteristics for the electrically initiated HBR elicited shorter onset latency, longer duration, and higher magnitude when the affected hand entered defensive peripersonal space (DPPS) for the face compared to when the hand was outside of it [47]. These findings were later reproduced by Bufacchi et al. [9-11], where they modeled the DPPS, and the factors that modulate its size. In a follow up study, they showed the attenuation of this effect when a screen was placed between the hand and the face, which was accompanied by a significant decrease in perceived threat ratings for the screen versus no screen conditions [45], suggesting that the screen either interrupted the integration of visual and tactile input, or as posited by the authors that the screen acted as a defensive shield against threats to the orofacial region. Finally, they showed that trigeminal neuralgia, a unilateral neuropathic pain disorder affecting the orofacial region, is associated with a unilateral expansion of the peripersonal space. Together, these data show that there is a unique DPPS for the orofacial region that can identify and integrate threats of potentially harmful stimuli towards the orofacial region.

Further evidence for the amplified perception of noxious stimuli in the orofacial region comes from studies that showed heat stimuli on the face are perceived as more painful than elsewhere on the body. Specifically, noxious heat delivered to the forehead was associated with higher fear of pain ratings and significantly higher within subject sensitization than the identical stimulus delivered to the volar forearm [50]. In a different study, a capsaicin-induced hyperalgesia applied to the face elicited increased ipsilateral amygdalar activity in response to thermal and brush stimuli on the sensitized side compared to the control side [39]. Cortical differences beyond the amygdala have been shown to differ between how orofacial versus bodily pain related fear modulate brain-wide activity, as well. Memory encoding for images associated with facial versus hand pain showed increased activity between the left amygdala and the visual processing, memory encoding and object recognition regions such as the left lateral occipital cortex and parahippocampal gyrus [49]. Object recognition was associated with increased connectivity between the left amygdala and the right parahippocampal gyrus in a different study that utilized the same experimental set up and stimuli [48]. The increased functional connectivity between the amygdala to memory regions has been speculated to account for higher fear elicited by the aversiveness and the potential threat of facial pain. Considered together, these findings suggest an enhanced amygdalar response to orofacial pain illustrated by increased physiological, functional connectivity and behavioral metrics in healthy individuals.

Chronic orofacial disorders are undergirded by pathologies of the trigeminal system, yet there exists a lack of clarity understanding the specific mechanisms that might produce enhanced negative affect present for trigeminal, as opposed to spinal, noxious input. Synthesizing the functional and behavioral findings from human subjects with the structural findings from rodent literature presents a challenge due to lack of translational studies providing structural targets to be used in studies with humans. We propose that future research on affective dimensions of orofacial pain can bridge this gap by studying function and structure in tandem across the novel trigeminal pathway shown in the current study.

### Limitations

Studying subcortical and brainstem structures *in vivo* has become more accessible due to increases in probabilistic atlases, gains in imaging resolution, and publicly available MRI data. However, compared to the degree of detail proffered by rodent tracing studies, circuit mapping in the brainstem-to-subcortical regions is inherently of coarser resolution. Indeed, in the current study, tractography was carried out at the gross CeA level, despite the animal literature indicating that the capsular and lateral, and not the medial subnuclei, receive parabrachial inputs. Similarly, we traced from the latPB, and not external latPB. To our knowledge, no probabilistic atlases to date provide seeds for the subnuclei for these two specific regions, and it is not clear that these could be reliably resolved with UHF MRI. We chose to assess unilateral direct CNV- latPB-CeA connectivity in consideration of the rodent literature. Given that the spinoparabrachial tract is comprised of both ipsilateral and contralateral efferents from PB [25,54], whether the novel CNV-latPB-CeA circuit in humans has a contralateral component remains an outstanding question. Although the ipsi-and contralateral components of the spinoparabrachial pathway have been shown to undergird different nociceptive processes in rodents at the spinal level [18], whether such differences are found for the trigeminal system has yet to be explored.

### Conclusion

Rodent work in the last decade has provided specific neural circuity undergirding affective aspects of nociceptive processing. We provide the successful resolution of the CNV-latPB-CeA white matter circuit *in vivo* for the first time. Future work involving human subjects can now investigate structural and functional differences along this pathway in experimental and clinical populations to further elucidate enhanced affect in response to orofacial pain.

## ACKNOWLEDGEMENTS

M Moayedi is supported by an NSERC Grant (RGPIN-2018-04908), the Bertha Rosenstadt Fund and the SEED program funded by the Dental Research Institute at University of Toronto’s Faculty of Dentistry. He also holds a Canada Research Chair (NSERC, Tier 2) in Pain NeuroImaging. Data were provided by the Human Connectome Project, WU-Minn Consortium (Principal Investigators: David Van Essen and Kamil Ugurbil; NIH grant #1U54MH091657) funded by the 16 NIH Institutes and Centers that support the NIH Blueprint for Neuroscience Research; and by the McDonnell Center for Systems Neuroscience at Washington University.

The authors have no conflicts to report.

