## Supplementary Figure 1 for "Delineation of the Trigeminal-Lateral Parabrachial-Central Amygdala Tract in Humans: An Ultra-High Field Diffusion MRI Study"

Short Title: CNV-latPB-CeA Connectivity

**Corresponding Author:**
Massieh Moayedi, PhD
Centre for Multimodal Sensorimotor and Pain Research
Faculty of Dentistry
University of Toronto
501B-123 Edward St
Toronto, ON
Canada M5G 1E2

**TABLE OF CONTENTS**

Supplemental Figures……………………………………………………………………………3

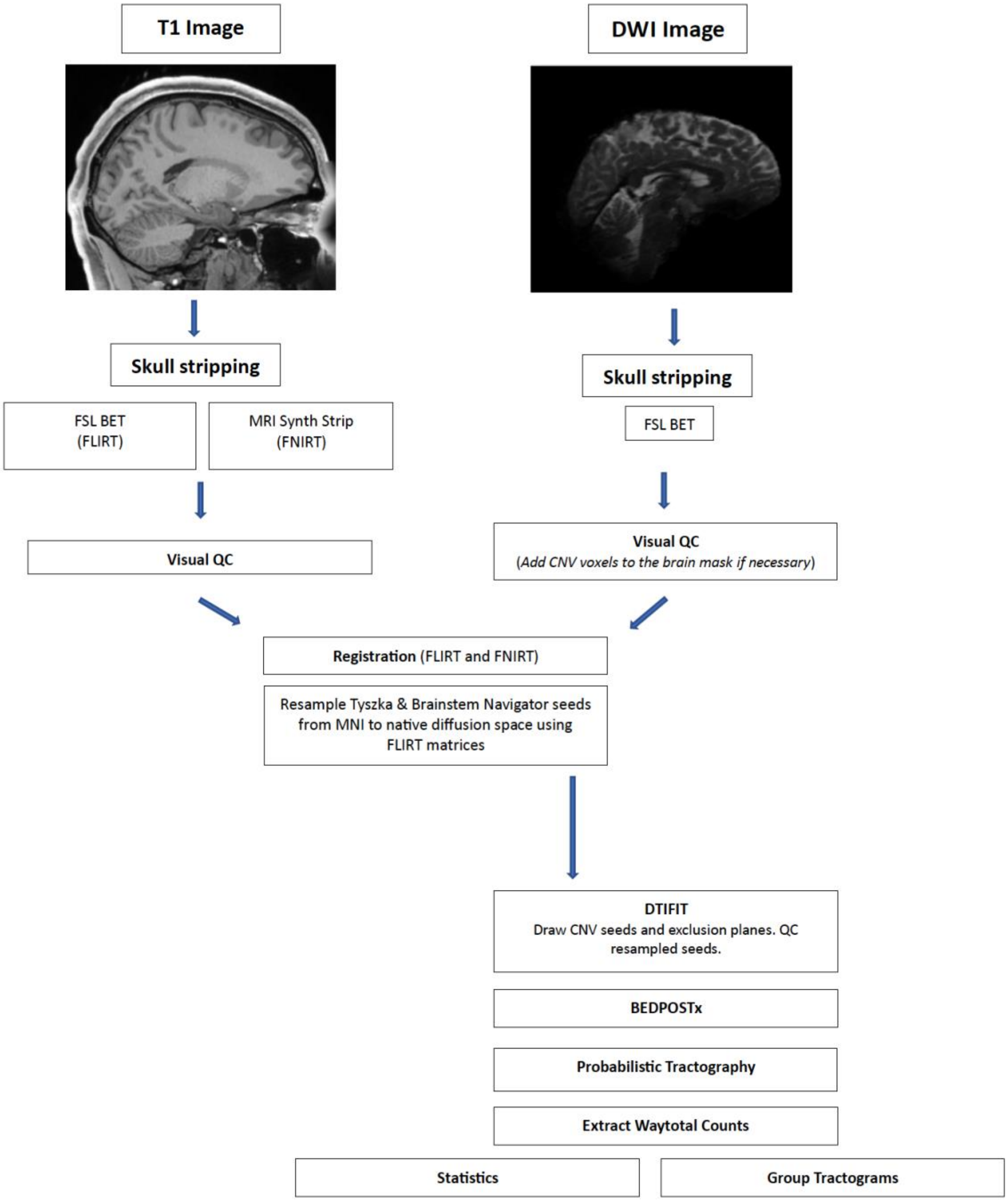


**Supplementary Figure 1: Study-specific preprocessing flowchart.**
Diffusion and T1-weighted structural images were registered to the MNI template brain using FLIRT and FNIRT algorithms. Probabilistic seeds in MNI space were resampled to native diffusion space using the FLIRT matrices to generate waytotal counts. Group tractograms were generated in two independent instances via the FLIRT and FNIRT transformations to allow for improved visualization of the trigeminal-lateral parabrachial-central amygdalar circuit.
